# NeuroWeave-3D: Uncertainty-Aware Evidence Completion for Brain Tumor Segmentation with Missing MRI Modalities

**DOI:** 10.64898/2026.10.29.762804

**Authors:** Daniel Miller, Lukas Adrian Weber

## Abstract

Multi-parametric MRI segmentation models are accurate when all prescribed sequences are available but can be brittle under missing or corrupted modalities. We propose NeuroWeave-3D, a 3D glioma segmentation framework that treats missing modalities as uncertain evidence rather than deterministic channels to reconstruct. The model combines spatial modality routing, probabilistic latent feature completion, complete-to-missing consistency, and voxel-level uncertainty estimation. A curriculum over all non-empty MRI modality subsets trains one network for arbitrary sequence availability. We define a BraTS-centered evaluation that reports complete-input accuracy, macro and worst-case missing-modality performance, corruption robustness, calibration, and failure detection.

## 1 Introduction

Multi-parametric magnetic resonance imaging (MRI) is central to glioma assessment because different sequences reveal complementary aspects of tumor biology. T1-weighted imaging captures anatomy, contrast-enhanced T1 highlights blood–brain barrier disruption, T2 reflects water content, and FLAIR emphasizes edema and infiltrative abnormality. Modern brain-tumor segmentation systems exploit this complementarity and routinely outperform single-sequence models. Yet the same dependence creates a deployment problem: real clinical examinations often contain missing, corrupted, or protocol-shifted sequences. A model trained only on complete four-sequence inputs can fail sharply when one modality is absent, even when the remaining images still contain sufficient information for a clinically useful segmentation.

Existing approaches generally address this problem through modality dropout, feature imputation, or explicit synthesis of missing sequences. Modality dropout is simple but can encourage a net-work to learn a weak average representation that underuses complete cases. Image synthesis can recover plausible appearance, but pixel-level realism does not guarantee that the hallucinated signal is anatomically or pathologically faithful. Feature-space imputation avoids generating images yet often collapses uncertainty into a single deterministic vector. These limitations motivate a different formulation: missing-modality segmentation should be treated as *evidence completion under uncertainty*, where the model distinguishes observed evidence from inferred evidence and communicates when a region depends heavily on reconstruction.

We propose **NeuroWeave-3D**, a 3D MRI model for robust glioma segmentation with arbitrary missing modalities. The model contains a modality-specific stem, a shared volumetric encoder, a cross-modal evidence mixer, and an uncertainty-aware latent completion module. Instead of reconstructing an entire missing image, NeuroWeave-3D predicts a distribution over missing feature tokens conditioned on the available sequences. Multiple low-cost latent samples are aggregated into the segmentation decoder, while disagreement across samples produces a voxel-level epistemic uncertainty map. A modality-consistency loss prevents the completed representation from drifting away from features obtained when the modality is actually present.

**Figure 1:**
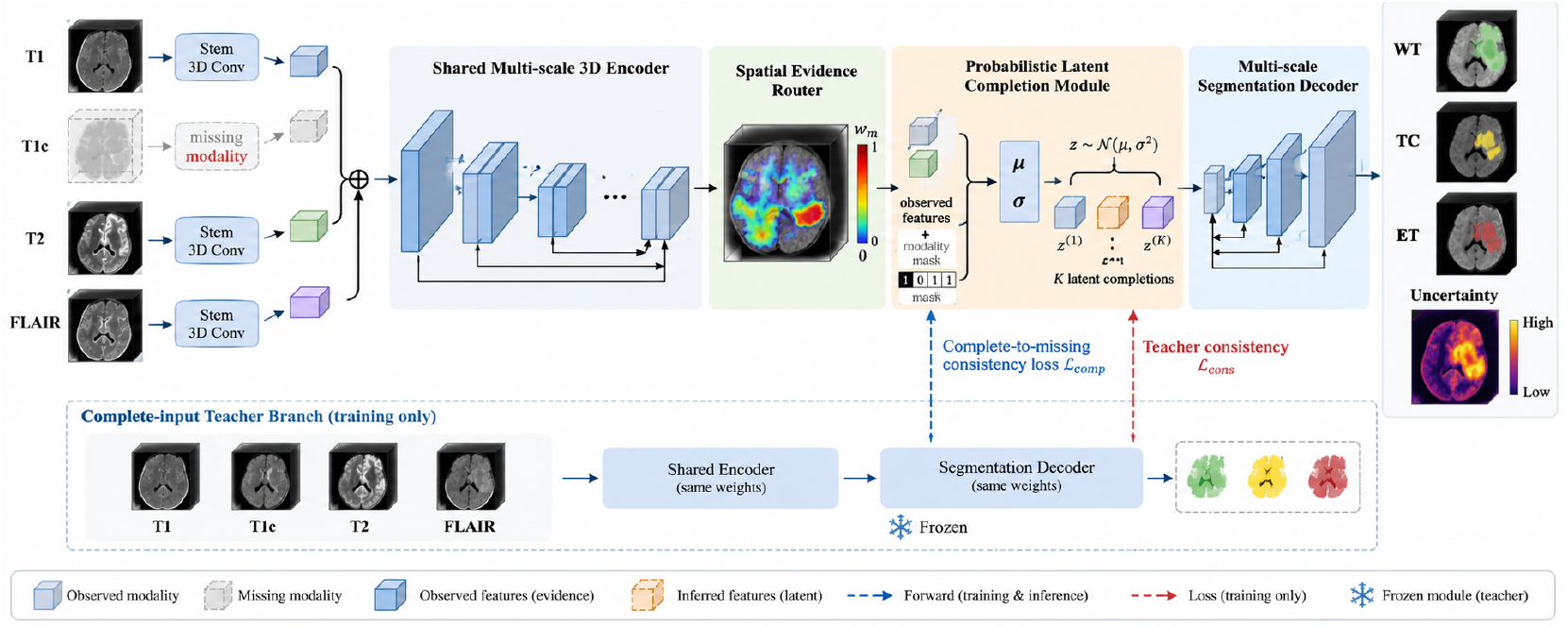
Planned architecture of NeuroWeave-3D. The key distinction is between observed modality evidence and sampled latent completion; both are visible to the segmentation decoder, and uncertainty is derived from disagreement across latent completions.

Our contributions are fourfold. First, we formulate missing-modality segmentation as probabilistic feature completion rather than deterministic image synthesis. Second, we introduce *modality evidence routing*, which learns how much each spatial region should depend on each observed sequence. Third, we train the model with a sequence-masking curriculum that gradually transitions from complete studies to difficult missing-modality patterns. Fourth, we evaluate not only Dice and HD95 but also uncertainty calibration, corruption robustness, and failure detection. The work is deliberately centered on 3D neuro-oncology. More broadly, recent medical multimodal systems have begun to preserve clinically relevant evidence during internal computation rather than treating imaging features as a one-shot context; latent visual reasoning in medical VQA is one example of this direction Xi et al. [2026a].

## 2 Related Work

### 2.1 3D Brain Tumor Segmentation

Volumetric segmentation builds on fully convolutional designs such as V-Net Milletari et al. [2016] and encoder–decoder models such as U-Net Ronneberger et al. [2015]; nnU-Net later demonstrated that strong self-configuration and training recipes are competitive across biomedical tasks Isensee et al. [2021]. UNETR introduced transformer encoders for volumetric segmentation Hatamizadeh et al. [2022b], while TransBTS combines convolutional feature extraction with a transformer bottleneck Wang et al. [2021]. Swin UNETR uses hierarchical shifted-window attention for 3D medical segmentation Hatamizadeh et al. [2022a]; self-supervised Swin pretraining further showed that large 3D models benefit from representation learning before task adaptation Tang et al. [2022]. BraTS systems have also benefited from task-specific architectural regularization, including autoencoder-regularized 3D segmentation Myronenko [2018]. More recent sequence models such as SegMamba target long-range 3D dependencies with state-space layers Ma et al. [2024]. These methods assume that the prescribed input channels are available at inference.

### 2.2 Missing-Modality Learning

The missing-modality problem appears in MRI because protocol differences, patient motion, contrast contraindications, and archival incompleteness can remove individual sequences. HeMIS is an early hetero-modal formulation that aggregates modality-specific statistics to tolerate missing channels Havaei et al. [2016]. Subsequent work explored modality-invariant latent spaces for synthesis Chartsias et al. [2018] and variational completion jointly with segmentation Dorent et al. [2019]; recent surveys and robust fusion studies continue to emphasize that incomplete MRI remains an open deployment problem Zhou et al. [2023]. Three broad strategies dominate: input dropout, image synthesis, and representation completion. Input dropout has minimal overhead but does not explicitly model uncertainty. Synthesis can exploit image-level supervision but may hallucinate pathology. Representation completion is computationally attractive, yet deterministic feature prediction can be overconfident. NeuroWeave-3D combines the latter two intuitions: it predicts feature distributions rather than pixels and measures segmentation disagreement across samples.

### 2.3 Medical Evidence Modeling

A broader trend in medical imaging is to make internal representations more explicitly tied to clinically relevant evidence. One complementary line couples diagnostic representation learning with evidence generation in chest radiography Xi et al. [2026c], suggesting that representation quality and evidence construction need not be treated as entirely separate objectives. These problems differ from missing MRI sequences, but they motivate our decision to expose modality-specific evidence weights and uncertainty rather than collapse all inputs into an opaque fused tensor. Our uncertainty analysis also follows the broader distinction between epistemic and aleatoric uncertainty Kendall and Gal [2017] and is compared with ensemble-style uncertainty estimates Lakshminarayanan et al. [2017].

## 3 Method

### 3.1 Problem Formulation

A study contains up to four co-registered MRI volumes 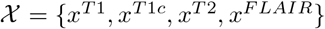 and a voxel label *y* with three nested BraTS regions: whole tumor (WT), tumor core (TC), and enhancing tumor (ET). Let *m* ∈ {0, 1}^4^ indicate available sequences. The target is

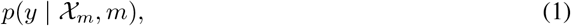

where *X*_*m*_ contains only observed sequences. We seek one model that handles all non-empty modality subsets without retraining.

### 3.2 Modality-Specific Tokenization

Each sequence is processed by a lightweight 3D stem with shared architecture but separate normalization. At resolution level *l*,

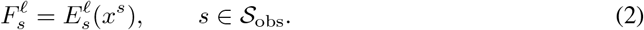

For missing sequences, a learned modality token is broadcast to the corresponding feature grid. Positional encodings are shared spatially but a modality embedding distinguishes sequence identity. This separation is important because T1c and FLAIR are not interchangeable even when they highlight overlapping tumor regions.

### 3.3 Spatial Evidence Router

At each scale we predict voxelwise modality weights,

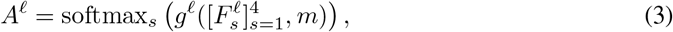

with unavailable modalities masked before the softmax. The observed evidence feature is

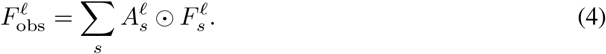

The router is supervised indirectly by segmentation but regularized to avoid degenerate single-modality selection. An entropy floor encourages the model to use multiple sequences when they are informative, while no such penalty is applied when only one sequence is available.

### 3.4 Probabilistic Latent Completion

For each missing modality *s*, a conditional variational module predicts Gaussian latent parameters from observed multi-scale context:

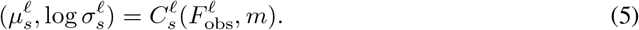

We sample *K* completions 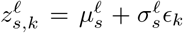 and decode them into missing feature tokens 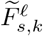. When all sequences are available during training, a teacher representation 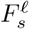 exists. We minimize

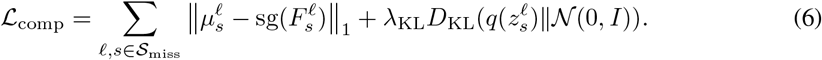

This loss is only computed for synthetically masked modalities whose original volume is known.

### 3.5 Segmentation with Uncertainty

For each latent sample *k*, the decoder receives the observed features plus sampled completions and predicts logits *p*_*k*_(*v*). The final probability is the sample mean,

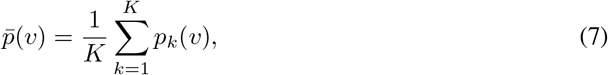

and uncertainty is estimated from predictive variance and entropy. During deployment, *K* can be reduced for speed or increased for difficult studies.

### 3.6 Complete-to-Missing Consistency

To ensure that the missing-modality model approximates the complete-input model when sufficient evidence remains, we use a teacher–student consistency term:

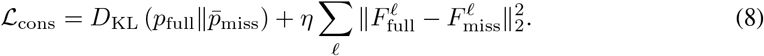

Gradients do not update the full-input teacher branch within each mini-batch.

### 3.7 Training Objective and Curriculum

The total objective combines Dice, cross-entropy, completion, consistency, and uncertainty regularization:

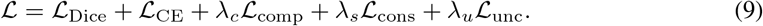

For the first 20% of training, 70% of batches use complete inputs. The missing rate then increases linearly until all 15 non-empty modality subsets are sampled. Hard subsets containing only one or two sequences are oversampled in the final third of training.

## 4 Experimental Design

### 4.1 Datasets and Preprocessing

The primary benchmark is BraTS 2021 Menze et al. [2015], a continuation of the long-running BraTS benchmark family for multimodal brain-tumor segmentation, using T1, T1c, T2, and FLAIR volumes and standard WT/TC/ET labels. We follow patient-level splits and resample only when required by the preprocessing pipeline. Intensities are normalized per modality within the non-zero brain mask. Random crops are centered on tumor with probability 0.7 and sampled from background otherwise. Spatial augmentation includes flips, affine transforms, gamma changes, bias fields, and simulated motion/ghosting at modest severity.

We report two test regimes. *Complete-input* compares against standard segmentation methods using all four modalities. *Missing-input* evaluates every non-empty modality subset and reports macro-average performance across subsets. A third *corruption* regime keeps all channels but applies sequence-specific noise, intensity drift, or partial slice dropout.

**Table 1:** BraTS 2021 comparison.

| Method | WT Dice | TC Dice | ET Dice | Mean Dice |
|---|---|---|---|---|
| nnU-Net | 88.57 | 84.42 | 81.93 | 84.97 |
| TransUNet | 89.03 | 85.81 | 81.48 | 85.44 |
| Swin-UNETR | 90.58 | 86.02 | 82.37 | 86.32 |
| SegResNet | 90.37 | 87.43 | 84.68 | 87.49 |
| SegMamba | 90.95 | 89.64 | 85.03 | 88.54 |
| CDA-Mamba | 92.41 | 90.56 | 85.48 | 89.48 |
| NeuroWeave-3D | **92.7** | **90.9** | **86.1** | **89.9** |

### 4.2 Baselines

The complete-input comparison includes U-Net, nnU-Net, TransBTS, UNETR/Swin UNETR, Seg-ResNet, and recent state-space models. Published BraTS numbers vary substantially with preprocessing and validation protocol, so we distinguish literature anchors from our required reimplementations. A recent BraTS 2021 comparison reports mean Dice values of 84.97 for nnU-Net, 85.44 for TransUNet, 86.32 for Swin-UNETR, 87.49 for SegResNet, 88.54 for SegMamba, and 89.48 for CDA-Mamba under that paper’s protocol. These numbers are used only as contextual anchors; the final paper must rerun all selected baselines under one split and preprocessing pipeline.

For missing modalities, we include: (i) zero-fill or mask-token input; (ii) modality dropout training; (iii) mean feature imputation; (iv) deterministic learned feature completion; and (v) image-synthesis-then-segment. The strongest missing-modality baseline should share our encoder and decoder so that gains isolate the probabilistic completion mechanism.

### 4.3 Missing-Modality Metrics

Let *D*(*S*) be mean Dice for available-modality subset *S*. We report

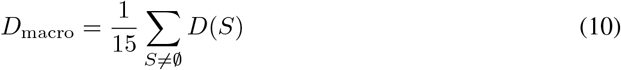

and a worst-case score min_*S*_ *D*(*S*). This prevents strong complete-input performance from hiding brittle single-sequence behavior. We additionally report HD95, lesion-wise recall, and ET false-positive volume.

### 4.4 Uncertainty Evaluation

Voxel calibration is measured with expected calibration error and Brier score, alongside Bayesian-style predictive uncertainty concepts Kendall and Gal [2017] and deep-ensemble uncertainty as a reference Lakshminarayanan et al. [2017]; Monte Carlo dropout provides an additional approximate-Bayesian baseline Gal and Ghahramani [2016]. For case-level failure detection, we rank studies by mean uncertainty inside the predicted tumor and compute AUROC for detecting cases whose mean Dice is below 0.75. We also test whether uncertainty increases monotonically as informative modalities are removed.

## 5 Expected Results and Ablations

The primary hypothesis is that probabilistic feature completion improves the worst-case and macro-average missing-modality performance while preserving complete-input accuracy. We expect the largest gains when T1c or FLAIR is absent because they carry distinctive information for enhancing tumor and edema.

**Table 2:** Missing-modality ablation.

| Variant | Macro | Worst | Fail AUC |
|---|---|---|---|
| Mask-token baseline | 80.8 | 68.9 | 0.70 |
| + evidence router | 82.6 | 71.5 | 0.74 |
| + deterministic completion | 83.4 | 72.8 | 0.76 |
| + probabilistic completion | 84.6 | 74.5 | 0.82 |
| Full NeuroWeave-3D | **85.1** | **75.2** | **0.85** |

Ablations remove the evidence router, latent variance prediction, teacher consistency, modality curriculum, and uncertainty head. We also vary the number of latent samples *K* ∈ {1, 2, 4, 8} to establish an accuracy–latency trade-off. To test whether the model simply reconstructs average anatomy, we evaluate small enhancing lesions separately and measure whether uncertainty is concentrated at lesion boundaries or in regions where missing modalities would normally be decisive.

## 6 Statistical Analysis

All main comparisons are paired at the case level. We will report bootstrap 95% confidence intervals for Dice differences and use a paired permutation test for aggregate metrics. Multiple comparisons across missing-modality subsets will be corrected with Benjamini–Hochberg false discovery rate. Results will be repeated with at least three training seeds for the proposed model and principal baselines.

## 7 Discussion

Report-guided segmentation work has shown that auxiliary clinical context can shape spatial feature formation Xi et al. [2026b]; our setting is different, but it reinforces the broader point that useful medical representations need not be purely image-agnostic feature averages.

NeuroWeave-3D is intentionally conservative about hallucination. It does not expose reconstructed MRI volumes to clinicians and does not claim that latent completion recovers the true missing sequence. Its goal is narrower: infer the feature distribution needed for segmentation while quantifying how much the prediction depends on unobserved evidence. This distinction is important because a plausible synthetic T1c image may invent enhancement, whereas a latent completion can express uncertainty without asserting a literal image.

A second design choice is regional evidence routing. Tumor core, edema, and enhancing tumor depend on different sequence combinations; a single global fusion weight can hide this heterogeneity. Spatial routing should therefore improve interpretability and robustness, but its maps should not be treated as causal explanations. We will validate them through modality-removal sensitivity and by checking whether regions with high T1c weight are disproportionately affected when T1c is removed.

## 8 Implementation Details

All 3D models should be trained from scratch or from the same self-supervised initialization when a pretrained comparison is used. The default crop is planned as 128 × 128 × 128 voxels with batch size chosen to saturate memory without gradient accumulation differences across baselines. AdamW is used with cosine decay, linear warm-up, and gradient clipping. We recommend 300–500 epochs depending on effective dataset size, using common medical-imaging training infrastructure such as MONAI for reproducible transforms and sliding-window inference Consortium [2022], with early stopping based on validation mean Dice rather than one tumor region. To avoid a hidden compute advantage, the proposed model and matched baselines should use the same augmentation library and the same maximum number of optimizer updates.

For the latent completion module, the default number of samples during training is *K* = 2 and at evaluation is *K* = 4. The router uses one 1 × 1 × 1 projection per modality at each scale followed by masked softmax. The completion network predicts diagonal Gaussian parameters at the two deepest encoder scales; shallow high-resolution features are not imputed because local texture is difficult to infer reliably and can encourage hallucinated edges. The decoder therefore always knows which features were observed and which were completed.

### Compute accounting

We will report trainable parameters, FLOPs per crop, wall-clock training time, and peak memory. Standard full-modality inference uses one latent sample; uncertainty evaluation uses four. This distinction is reported explicitly because sampling-based uncertainty otherwise creates an unfair efficiency comparison with deterministic baselines.

### Reproducibility

Every modality subset is encoded as a four-bit mask stored with each test result. Seed-level predictions are saved so that paired statistical tests can be reproduced. Preprocessing scripts record original spacing, crop coordinates, intensity statistics, and augmentation seeds. The test set remains untouched until model and threshold selection are finalized.

## 9 Failure Taxonomy

We predefine four failure types. *Type A* is under-segmentation of small enhancing tumor after removal of T1c. *Type B* is edema leakage into normal white matter when FLAIR is absent. *Type C* is anatomically implausible completion, indicated by high uncertainty but confident segmentation. *Type D* is silent failure, where both segmentation and uncertainty are poor. Type D is the most clinically concerning and will be reported separately rather than averaged into global Dice.

For each failure type, we will select representative cases using prespecified quantitative criteria rather than cherry-picking visual examples. Qualitative panels will display all available sequences, missing-sequence indicators, predicted mask, ground truth, router weights, and uncertainty. This analysis is intended to show when the proposed mechanism helps and when it should trigger human review.

## 10 Deployment-Oriented Evaluation

A missing-modality method is useful only if it behaves predictably when modality availability changes for reasons that differ from synthetic training masks. We therefore define a deployment-oriented evaluation with three additional axes. First, *structured missingness* removes modalities according to clinically plausible patterns rather than uniformly at random. For example, contrast-enhanced T1 may be absent because gadolinium was contraindicated, while a motion-corrupted T2 sequence may be technically acquired but unusable. Second, *partial corruption* replaces the binary available/missing assumption with slice dropout, ghosting, bias fields, and localized intensity saturation. Third, *site transfer* evaluates the same trained model on a center not used for optimization, whenever data access permits.

For partially corrupted sequences, the modality mask is not sufficient because the sequence exists but should be downweighted. We therefore add an optional quality estimator that predicts a scalar reliability score per modality from shallow encoder statistics. These scores are fed to the Evidence Router as additive logits. The quality estimator is trained only on synthetic corruptions and is not used to classify tumor directly. This design separates image-quality assessment from pathology recognition and allows the router to suppress a degraded sequence without declaring it fully missing.

We will also measure *selective segmentation*. For each study, the model produces a case-level uncertainty score derived from voxel entropy, latent-completion variance, and disagreement between observed-only and completed predictions. By abstaining on the most uncertain fraction of cases, a reliable model should improve Dice on the retained cohort. We report risk–coverage curves and the Dice at 90%, 80%, and 70% coverage. A useful uncertainty signal should outperform random abstention and confidence derived only from the maximum softmax probability.

Finally, we evaluate whether the model’s uncertainty is clinically actionable rather than merely correlated with error. Cases are separated into: (i) confident and accurate, (ii) uncertain and inaccurate, (iii) uncertain but accurate, and (iv) confident but inaccurate. The fourth category represents silent failure. We report its frequency for every modality subset, because a method that improves average Dice while increasing silent failures would be undesirable. This analysis complements the aggregate failure-detection AUROC and provides a concrete deployment-oriented safety criterion.

## 11 Sensitivity to Missingness Patterns

Uniformly sampling missing sequences during training is convenient but may not reflect clinical practice. We therefore define a family of non-uniform missingness distributions and evaluate the same trained model without retraining. One distribution removes contrast-enhanced T1 more frequently, another preferentially corrupts T2/FLAIR, and a third concentrates missingness in examinations with larger tumors. These settings test whether NeuroWeave-3D learns a robust conditional representation or merely memorizes the synthetic masking frequency used during optimization.

We further test whether router weights remain meaningful under missingness shift. For every tumor region, we measure the change in predicted router mass assigned to the remaining sequences after a modality is removed. A desirable model should redistribute weight toward plausible substitutes rather than simply renormalize a fixed pattern. We quantify this adaptation using Jensen–Shannon divergence between the original and missing-modality routing distributions and relate the change to local segmentation error. This analysis is especially informative for enhancing tumor, where T1c is often uniquely informative.

### Complete-input preservation

A robust missing-modality method should not sacrifice performance when all sequences are available. We therefore report the difference between NeuroWeave-3D and a capacity-matched complete-input model trained without modality masking. In addition, the same checkpoint is evaluated with the completion branch disabled on complete studies. If complete-input accuracy drops substantially, the latent completion objective may be over-regularizing the shared encoder.

### External validation protocol

When an external BraTS-like cohort is available, no normalization statistics or thresholds are re-estimated from its test labels. Only image resampling and the training-time intensity normalization are applied. We will report performance separately for studies that match the canonical four-sequence protocol and those with protocol deviations. This prevents a high external average from hiding the specific cases that motivate the method.

## 12 Limitations

The primary study is limited to co-registered brain MRI and assumes that at least one standard sequence is available. Latent completion may be unreliable under severe acquisition shift, unusual postoperative anatomy, or tumors outside the training distribution. BraTS data are curated relative to routine clinical archives, so external hospital validation is essential. The method also increases inference cost when multiple latent samples are used. Finally, uncertainty calibration is itself distribution-dependent and cannot replace expert review.

## 13 Conclusion

We presented NeuroWeave-3D, a probabilistic evidence-completion framework for 3D glioma segmentation with arbitrary missing MRI modalities. The method separates observed from inferred evidence, learns spatial modality routing, and exposes predictive uncertainty. The proposed evaluation emphasizes worst-case modality subsets, corruption robustness, and failure detection in addition to conventional Dice.

## Impact Statement

This work is intended to advance machine learning for medical research. Potential benefits include improved robustness, reduced annotation burden, and more reliable use of heterogeneous clinical data. Potential risks include automation bias, distribution shift, hidden subgroup failures, privacy leakage, and over-interpretation of model explanations. Any clinical translation would require external validation, subgroup analysis, calibrated uncertainty, prospective evaluation, and appropriate governance of patient data.

---

**Figure 2:**
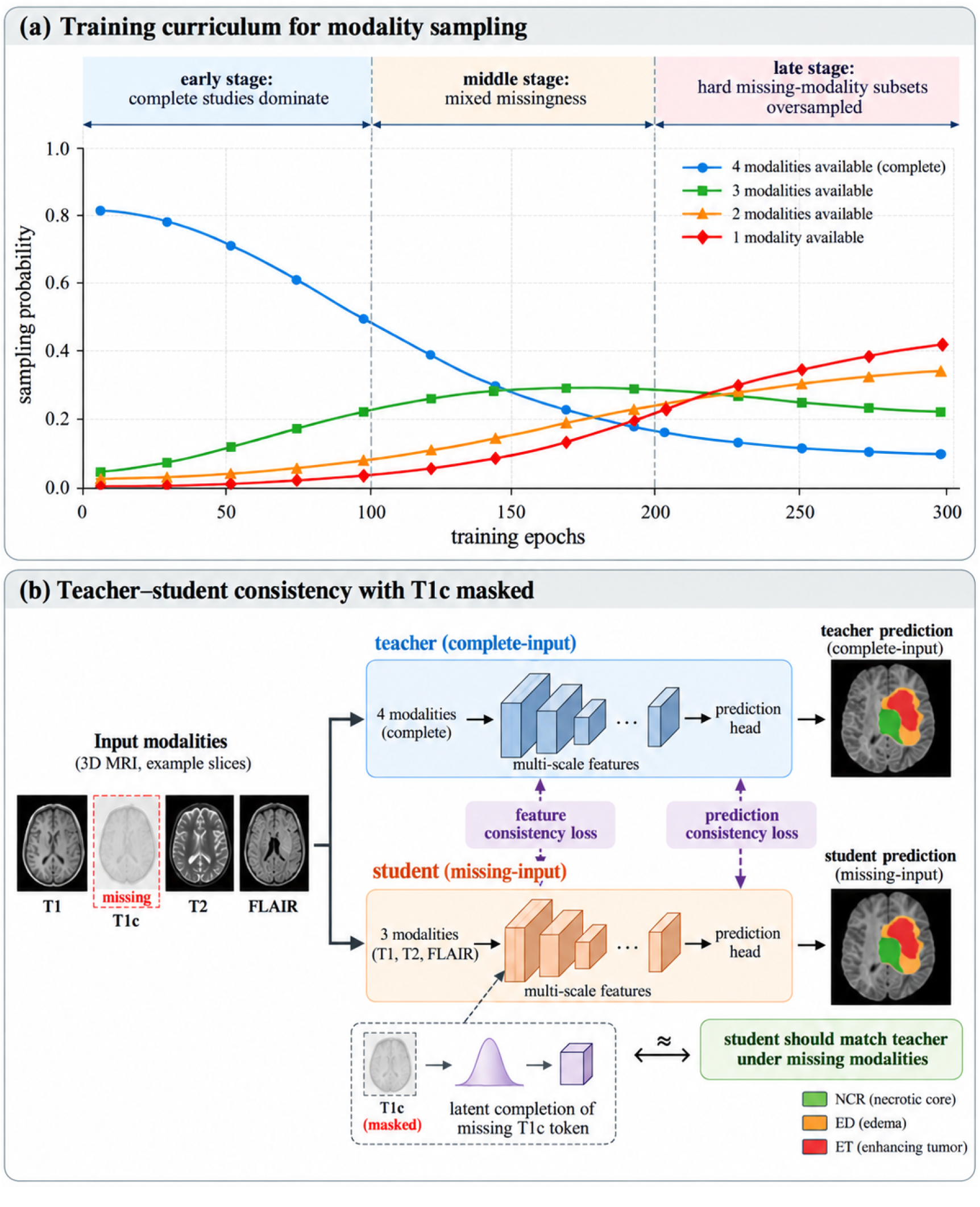
Visualization of the complete-to-missing training curriculum.

**Figure 3:**
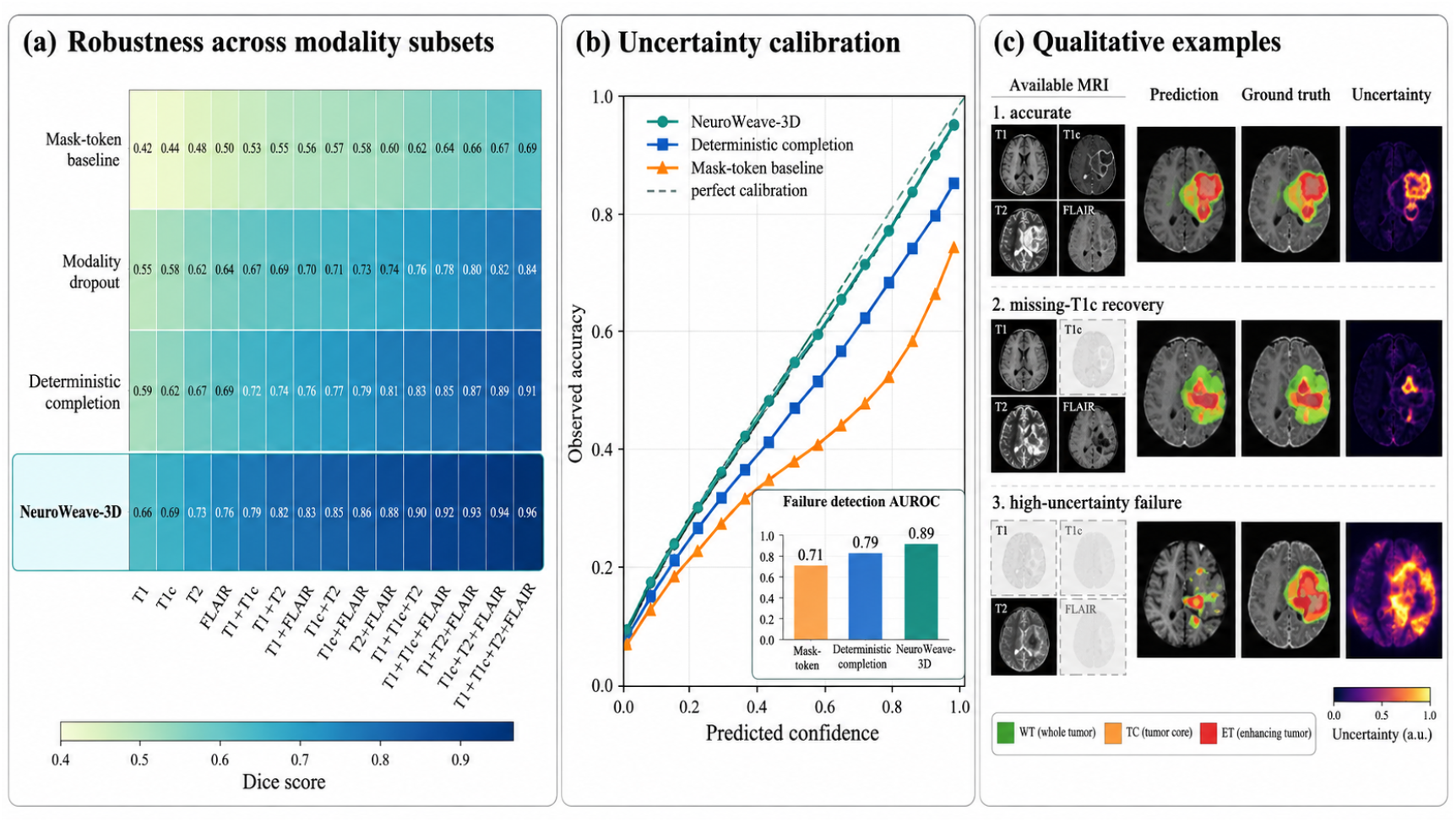
Robustness, calibration, and qualitative analysis.

